# Seed-competent α-synuclein pathology in metachromatic leukodystrophy: the expanding spectrum of α-synucleinopathy in sphingolipidoses

**DOI:** 10.1101/2024.08.09.607301

**Authors:** Simona Ghanem, Jade Hawksworth, Searlait Thom, Annabelle E. Hartanto, Joseph O’Neill, Janarthanan Ponraj, Said Mansour, Johannes Attems, Angela Pyle, Lauren Johnson, Jack Baines, Robert W. Taylor, Tiago F. Outeiro, Omar M.A. El-Agnaf, Daniel Erskine

## Abstract

Metachromatic leukodystrophy (MLD) is a rare - typically paediatric - sphingolipid storage disorder resulting from bi-allelic pathogenic variants in the *ARSA* gene, encoding the lysosomal arylsulphatase A (ASA). Heterozygous variants in *ARSA* are associated with risk of Lewy body diseases (LBD), a group of age-associated neurodegenerative disorders characterised by the accumulation of the protein α-synuclein; however, no study has yet determined whether α-synuclein with putative pathological features is observed in MLD brain tissue. We examined *post-mortem* brain tissue from MLD cases (N=5, age 2-33) compared to matched control cases using histological approaches and α-synuclein seeding amplification assay (SAA). Juvenile-onset MLD cases exhibited granular α-synuclein deposits in neurons of regions prone to neuronal pathology in MLD, and seed-competent conformers that generated atypical short, twisted fibrils on SAA. In contrast, infantile-onset MLD cases gave only variably positive reactions on SAA. In summary, this study suggests MLD cases manifest α-synuclein pathology reminiscent of that observed in LBD, even in juvenile populations, further expanding the spectrum of sphingolipid storage disorders associated with the aggregation of α-synuclein. These findings have important implications for understanding the disease process of both LBD and MLD, potentially highlighting novel pathways for therapeutic interventions in both conditions.

## MAIN TEXT

Lewy body diseases (LBD), such as Parkinson’s disease (PD) and dementia with Lewy bodies, are collectively the second most common neurodegenerative disorder after Alzheimer’s disease [1]. Neuropathologically, LBD are characterised by deposits of the protein α-synuclein as spherical intracellular deposits termed Lewy bodies that form in vulnerable populations of neurons throughout the brain [2]. Although the direct relevance of Lewy bodies to the neurodegeneration that underlies the clinical phenotype of LBD is currently unknown, the aggregation and deposition of α-synuclein is thought to be a key pathogenic event in LBD, and remains a major target for candidate disease-modifying therapeutics [3]. However, despite the apparent importance of the aggregation of α-synuclein in LBD, the factors that underlie it remain largely unknown.

A number of LBD risk genes encode enzymes that catabolise sphingolipids, a group of structural and signalling lipids, and include genes such as *GBA1, GALC*, and *SMPD1* [4-7]. We previously reported the presence of α-synuclein aggregates reminiscent of those observed in LBD in *post-mortem* brain tissue of infants with Krabbe disease [9]. The identification of a neuropathological feature associated with increased age in the brains of infants is notable, as it suggests a direct link between the aetiology of Krabbe disease and α-synuclein aggregation. Such information is important as despite the central role ascribed to α-synuclein aggregation in LBD, the mechanisms underlying its occurrence remain unknown. Sphingolipidoses have a monogenic aetiology that is typically well-defined; therefore, they may provide important insights into understanding factors that contribute to α-synuclein aggregation more broadly.

MLD is a rare lysosomal storage disorder typically caused by pathogenic bi-allelic variants in the *ARSA* gene, which encodes a lysosomal enzyme, arylsulphatase A (ASA), responsible for the de-sulphation of sulphatide/sulpho-galactosylceramide [10]. ASA dysfunction leads to the accumulation of its substrate, sulphatide, which is thought to underlie progressive loss of white matter that characterises MLD [10]. *ARSA* has recently been identified as an LBD risk gene, the ASA substrate sulphatide is elevated in LBD brain tissue, and a previous study indicated knockdown of *ARSA* increases the abundance of α-synuclein *in vitro* [6, 11-13]. One previous study has examined α-synuclein histologically in MLD brain tissue, reporting glial immunoreactivity in the putamen with a pan-α-synuclein antibody in two MLD cases [14]. However, it is not clear if such α-synuclein pathology manifests putative pathogenic attributes thought to be associated with LBD, such as phosphorylation at serine 129 or seed-competence on the α-synuclein seeding amplification assay (SAA); thus, the extent to which the disease process of MLD mirrors that in LBD remains undetermined. Therefore, we sought to identify whether α-synuclein depositions are observed in the MLD brain, and to determine whether α-synuclein in MLD brain tissue is seed-competent.

Ethical approval for this study was granted by Newcastle University Ethical Review Board (Ref: 18241/2021). Tissue from MLD cases (N=5) and age-, sex- and ethnicity-matched controls (N=5) were obtained from the University of Maryland Brain and Tissue Bank through the NIH NeuroBioBank (Table 1). To confirm pathogenic variants in *ARSA* in the MLD cohort, DNA was extracted from the cerebellar white matter using DNeasy Blood and Tissue DNA purification kits (Qiagen). Primer pairs flanking the variant of interest in *ARSA* (NM_000487.6) were custom designed using primer3 (http://primer3.ut.ee/). Patient DNA was PCR-amplified using GoTaq G2 HotStart Polymerase (Promega). PCR products were purified using Exonuclease I and FastAP (ThermoFisher Scientific) and then Sanger sequenced (BigDye® Terminator v3.1 Sequencing Kit, ThermoFisher Scientific) on an Applied Biosystems 3500XL Genetic Analyzer (ThermoFisher Scientific). Data were analysed using SeqScape v4 (Applied Biosystems). Pathogenicity scores were ascribed based on the classification system outlined in {Richards, 2015 #243}. All samples manifested bi-allelic variants in *ARSA*, both juvenile-onset MLD cases, MLD 4 and 5, manifested the most common ARSA variant associated with MLD, c.1283C>T [16]. Three cases manifested variants in the second most common c.465+1G>A, MLD 1, 3 and 5, of which MLD 3 was in a homozygous state (Table 1).

**Table 1:** Demographic details of the cohort used in the present study.

| Case ID | Diagnosis | Age at death (years) | Sex | Ethnicity | Genetic diagnosis | Cause of death | Dentate | Globus pallidus |
| --- | --- | --- | --- | --- | --- | --- | --- | --- |
| *MLD* |  |  |  |  |  |  |  |  |
| MLD 1 | MLD | 2 | F | Black | c.1130_1132delTCT / c.465+1G>A | Complications of disorder | X |  |
| MLD 2 | MLD | 3 | M | White | c.847G>T / c.1010A>T | Complications of disorder | X | X |
| MLD 3 | MLD | 6 | M | White | c.465+1G>A (Hom) | Complications of disorder | X | X |
| MLD 4 | MLD | 17 | F | White | c.1283C>T (Hom) | Complications of disorder | X | X |
| MLD 5 | MLD | 33 | M | Black | c.465+1G>A / c.1283C>T | Complications of disorder | X | X |
| *Controls* |  |  |  |  |  |  |  |  |
| C 1 | Control | 2 | F | Black |  | Acute myocarditis |  | X |
| C 2 | Control | 2 | F | Black |  | Drowning | X | X |
| C 3 | Control | 3 | M | White |  | Ventricular septal defect |  | X |
| C 4 | Control | 3 | M | White |  | Blunt force injuries of the head and torso | X |  |
| C 5 | Control | 4 | M | White |  | Head and neck injuries | X |  |
| C 6 | Control | 6 | M | White |  | Multiple injuries from a motor vehicle accident |  | X |
| C 7 | Control | 17 | F | White |  | Multiple injuries |  | X |
| C 8 | Control | 33 | M | Black |  | Fentanyl intoxication and atherosclerotic CVD | X | X |

To characterise neurodegenerative pathology in MLD cases, 5μm sections of formalin-fixed paraffin-embedded tissue from the dentate nucleus and globus pallidus were dewaxed in Histoclear and rehydrated through a graded series of ethanol solutions until water. Heat-mediated antigen retrieval in citrate buffer pH6 was performed to unmask epitopes, followed by immersion in formic acid for 5 minutes, prior to blocking in 10% normal goat serum and incubation in primary antibodies (anti-pS129, ab51253, Abcam, Cambridge, UK; anti-4G8, #800708, BioLegend, CA, USA; anti-AT8, #MN1020, Invitrogen, MA, USA). Sections were incubated in primary antibodies for one hour at room temperature prior to washing and probing with the universal probe and polymer-HRP components of the Mach 4 Polymer kits, as per manufacturer’s instructions (CellPath, PBC-M4U534L). Sections were imaged on an AxioImager microscope (Zeiss). pS129 immunoreactivity showed a relatively consistent picture of granular intraneuronal accumulation of α-synuclein pS129 in neurons of the dentate nucleus and globus pallidus in juvenile-onset MLD cases (cases 4 & 5, aged 17 and 33 years old, respectively; Figure 1C & F). Additionally, case 4 also manifested pS129-immunoreactive spherical structures in the globus pallidus that resembled axonal spheroids (Figure 1F). There was no histological evidence of α-synuclein deposition in infantile MLD cases or controls. Amyloid-β staining was similarly observed as intraneuronal deposits in juvenile-onset MLD cases and, to a lesser extent in some infantile-onset cases, but no extracellular plaques were observed in any case (Supplementary Figure 1). Tau pathology was not observed in any region of any case.

**Figure 1:**
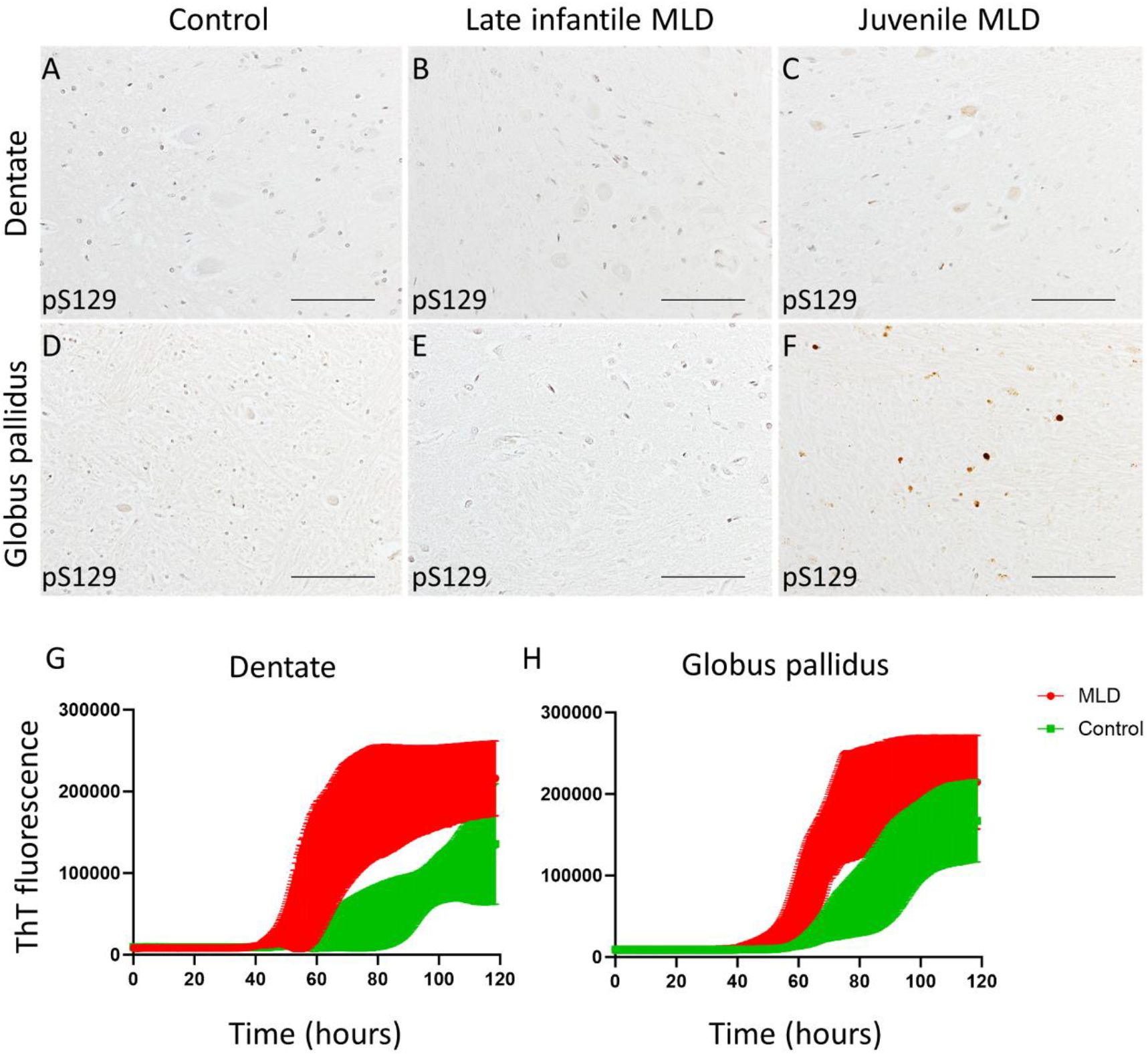
α-synuclein pathology in MLD cases. pS129 revealed minimal staining in the dentate nucleus of control and infantile MLD cases (A, B, D, E); whereas, granular intracellular deposits were observed in globus pallidus and dentate nucleus of both juvenile-onset MLD cases (C & F). SAA analysis identified significant differences at the group level between MLD and control cases in the dentate nucleus (G) and globus pallidus (H). Scale bars = 100 μm.

To determine the extent to which α-synuclein in MLD has qualities associated with LBD, we next performed α-synuclein seeded amplification assay (SAA) on the TBS-soluble fraction of tissue lysates from the globus pallidus and dentate nucleus of MLD and control cases. The SAA reaction buffer was composed of 0.1 M piperazine-N,N′-bis (ethanesulfonic acid) (Pipes; pH 6.9), 0.1 mg/mL recombinant α-syn and 10 μM thioflavin-T (Th-T). Reactions were performed in triplicates in a black 96-well microplate with a clear bottom (Nunc) with 85 μL of the reaction mix loaded into each well together with 15 μL of 0.1 mg/mL TBS-soluble fractions. The plate was then sealed with a sealing tape (ThermoFisher) and incubated at 37 °C for 120 h in a BMG FLUOstar OMEGA plate reader with intermittent cycles of 1-min shaking (500 rpm, double orbital) and 15-min rest throughout the indicated incubation time. Th-T fluorescence measurement, expressed as arbitrary relative fluorescence units (RFU), was taken with a bottom read every 15 min using 450 ± 10-nm (excitation) and 480 ± 10-nm (emission) wavelengths. The relative seeding activities of the assayed samples were presented by graphing fluorescence readouts against assay time. Statistical analysis was performed in GraphPad Prism v10.1.2. To compare SAA data, simple linear regression analysis was performed on the generated curves, from which direct comparison of slope and intercept was performed. The alpha value was 0.05 for all analyses. SAA analysis of tissue lysates from MLD cases identified a curve that differed from controls in both the dentate nucleus (F=1296, p<0.0001) and globus pallidus (F=554.6, p<0.0001; Figure 1).

Due to the phenotypic and pathological differences between late infantile and juvenile -onset MLD cases, comparisons were also made between these and their matched controls separately. Comparison of only late infantile cases and matched controls demonstrated differences between MLD cases and controls in dentate nucleus (F=1117, p<0.0001) but not the globus pallidus (p=0.3443; Figure 2 A&B). Juvenile-onset MLD cases manifested differences from controls in SAA reaction in both dentate nucleus (F=844.1, p<0.0001) and globus pallidus (F=1999, p=<0.0001; Figure 2 C&D).

**Figure 2:**
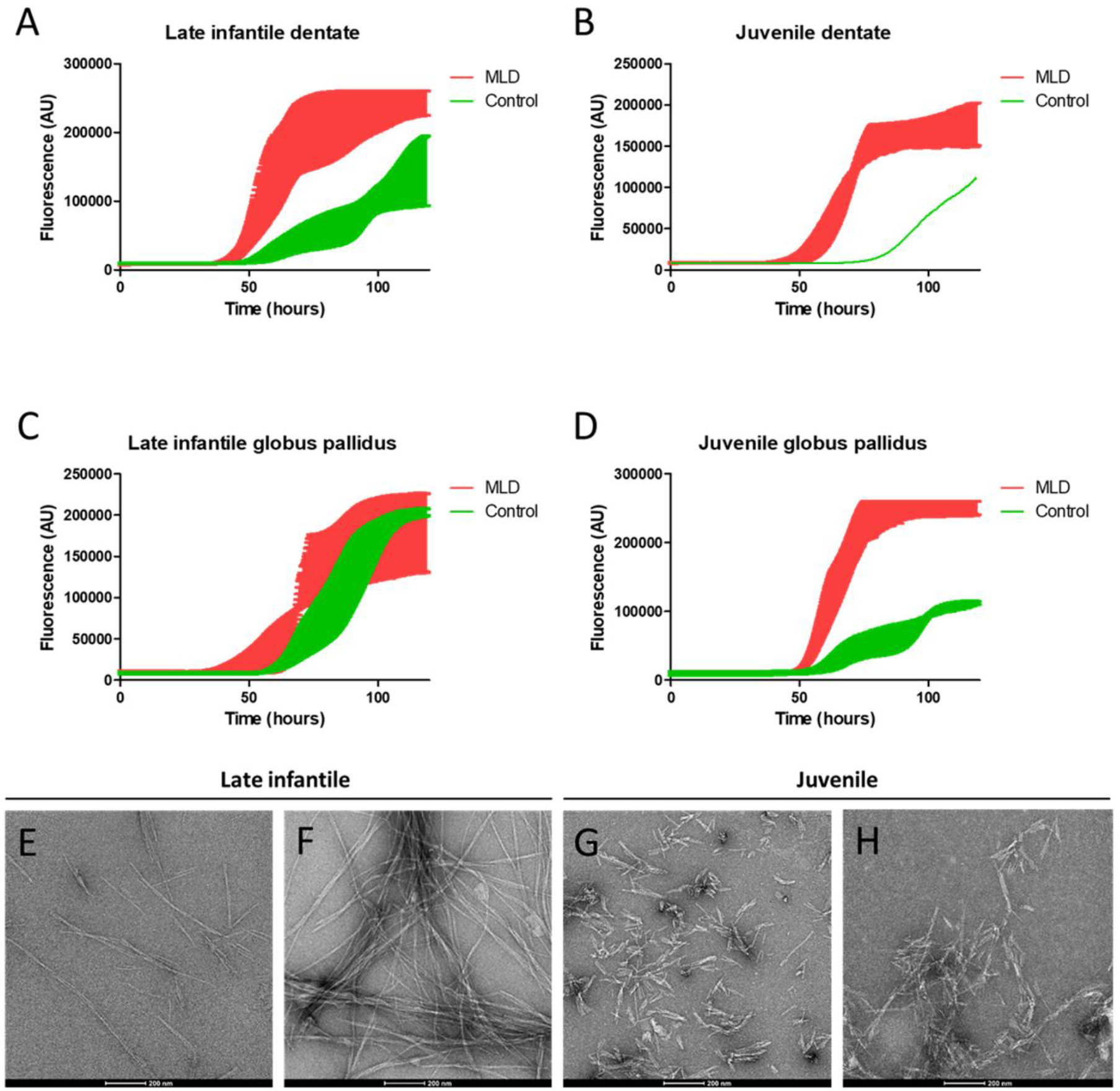
MLD samples manifest seed-competent α-synuclein conformers. SAA of dentate nucleus tissues revealed both late infantile and juvenile -onset MLD cases showed clear separation from control cases (A-B). In the globus pallidus, there was no clear separation between late infantile MLD cases and controls; in contrast, juvenile-onset MLD cases showed clear separation from controls (D). Electron microscopy of SAA end-products showed fine elongated fibrils generated from late-infantile cases (E-F); however, end-products from juvenile-onset MLD cases had an atypical twisted morphology and shorter length (G-H). Scale bars = 200 nm (E-H).

To confirm the presence of fibrils in end-products from MLD cases, SAA end-products were transferred in a 5 μl volume to a transmission electron microscopy grid (S162 Formvar/Carbon, 200 Mesh, Agar Scientific). After 1 min, samples were fixed using 5 μl of 0.5% glutaraldehyde followed by one wash with 50 μl of deionized water. Then 5 μl of 2% uranyl acetate were added to the grid, and after 2 min uranyl acetate was blotted off and grids were dried before placing sample holders before visualisation. Electron microscopy of SAA end-products from MLD cases demonstrated characteristic fibrillar structures (Figure 2E-H). However, it was notable that clear morphological differences were present in late-infantile MLD cases compared to those of juvenile onset. In particular, end-products of both juvenile-onset MLD cases manifested short fibrils of approximately 100-200 nm in length (Figure 2G & H), in contrast with fibrils exceeding 500 nm in late-infantile cases (Figure 2E & F).

This study reports, for the first time, that brain tissue from MLD cases contains seed-competent α-synuclein conformers. In particular, juvenile-onset MLD cases had histological evidence of intraneuronal α-synuclein aggregation, clear evidence of seed-competent α-synuclein conformers in two brain regions, and morphologically distinct fibrils generated from SAA. In contrast, late-infantile cases demonstrated seed competence on SAA, but this was not consistent across brain regions or cases. Taken together, these findings add to the growing spectrum of neurometabolic diseases caused by bi-allelic variants in genes associated with LBD that manifest pathological changes to α-synuclein, even in paediatric populations. The identification of α-synuclein pathology in monogenic disease cases much younger than the age at which α-synuclein typically sporadically develops suggests aetiological links between these diseases and α-synuclein aggregation. Therefore, these findings provide unique insights into mechanisms underlying α-synuclein aggregation that could be relevant to LBD and other common synucleinopathies.

Seed-competent α-synuclein pathology was not a universal observation across all regions of MLD cases; MLD 2 and 3 did not give consistent results across both regions, and manifested seed-competent α-synuclein pathology in only one region. In contrast, both juvenile-onset MLD cases, MLD 4 and 5, manifested both histological evidence of α-synuclein deposition and positive reactions on α-synuclein SAA. It is not clear why α-synuclein pathology is more consistent across regions in juvenile-onset MLD; however, one could speculate that the longer disease duration of juvenile-onset MLD is more permissive to the development of α-synuclein pathology than the relatively rapid course of late infantile MLD. It is notable that both juvenile-onset MLD cases had received bone marrow transplants during life though it is not clear how this would impact α-synuclein aggregation propensity beyond prolonging life to enable α-synuclein aggregation to occur. A previous murine study using models of Krabbe disease, a related sphingolipid storage disorder associated with α-synuclein pathology, reported that bone marrow transplant had no impact on cerebral α-synuclein deposition [21]. Nevertheless, it is not possible to rule out a potential uncharacterised impact of bone marrow transplantation on α-synuclein aggregation.

The present findings indicate that MLD cases manifest changes to α-synuclein reminiscent of those observed in LBD, with implications for the development of novel therapies for both conditions. Considerable drug discovery efforts are currently dedicated to therapies targeting α-synuclein aggregation in LBD, and the present findings could indicate repurposing these therapies for MLD could be a useful strategy [18]. The presence of α-synuclein pathology, a feature normally associated with advanced age, in cases aged between 2 and 33 years old, suggests a direct association between the metabolic dysfunction associated with MLD and, thus, may provide insights into factors that underlie α-synuclein aggregation in LBD. Furthermore, this could indicate that therapies in development to target the metabolic dysfunction in MLD, such as substrate reduction therapy, could be effective strategies to attenuate α-synuclein aggregation in LBD [19]. Similar approaches have already been employed for related metabolic pathways in LBD, such as glucocerebrosidase [20].

These findings add to the growing spectrum of rare neurometabolic diseases associated with seed-competent α-synuclein conformers, with implications for understanding the biology that underlies this pathological feature in diseases such as LBD. Furthermore, given the central role ascribed to α-synuclein aggregation in LBD pathogenesis, these findings may also indicate this process is implicated in MLD and a novel target for future therapeutics in this condition. Further studies are warranted to determine the role of α-synuclein pathology in MLD but its presence in non-LBD cases is important in the context of current biological staging schemes, which classify cases as having synucleinopathy or even Parkinson’s disease primarily on the basis of α-synuclein SAA [24, 25]. The present observations should prompt discussions of how these classification schemes can accommodate SAA-positive α-synuclein pathology in cases where the underlying molecular aetiology is well-characterised and accepted to result from other factors.

In summary, the present study has identified seed-competent α-synuclein pathology in brain tissue of MLD cases. These findings add to the growing spectrum of neurometabolic diseases associated with α-synuclein pathology, with important implications for understanding biological factors underlying LBD and the classification of cases based on SAA-positive α-synuclein pathology.

## ACKNOWLEDGEMENTS

Funding for this study was received by the Alzheimer’s Research UK Northern Network. DE is funded by an Alzheimer’s Research UK Senior Fellowship (ARUK-SRF2022A-006).

RWT is funded by the Wellcome Centre for Mitochondrial Research (203105/Z/16/Z), the Mitochondrial Disease Patient Cohort (UK) (G0800674), the Medical Research Council (MR/W019027/1), the Lily Foundation, Mito Foundation, the Pathological Society, the UK NIHR Biomedical Research Centre for Ageing and Age-related disease award to the Newcastle upon Tyne Foundation Hospitals NHS Trust, LifeArc and the UK NHS Highly Specialised Service for Rare Mitochondrial Disorders of Adults and Children.

