## Supplementary Figure 1 for "Seed-competent α-synuclein pathology in metachromatic leukodystrophy: the expanding spectrum of α-synucleinopathy in sphingolipidoses"

Supplementary materials

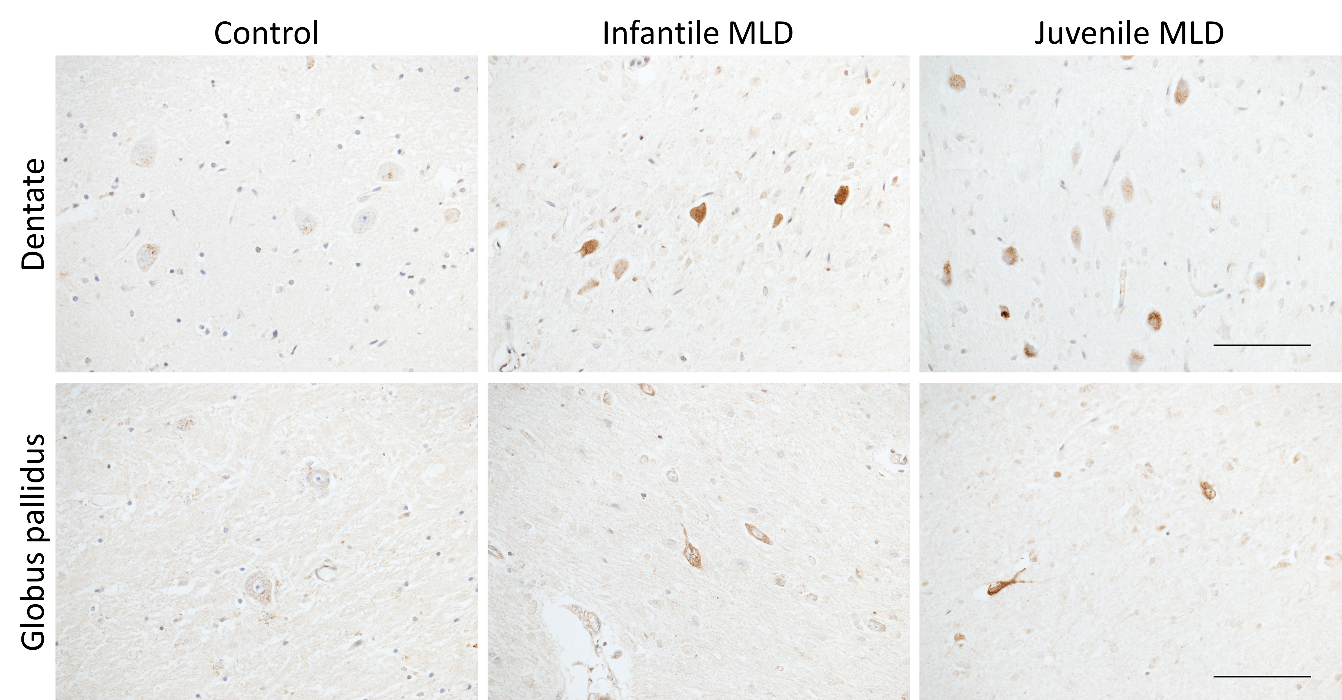
Supplementary Figure 1: 4G8 immunoreactivity in the dentate nucleus and globus pallidus of control and MLD cases. Amyloid-β immunoreactivity was observed in the cytoplasm of neurons of both regions, but was especially prominent in juvenile-onset MLD cases, but no extracellular plaques were observed. Scale bars = 100 µm.
